# Systematic Mining of gut microbiota biomarkers for IBD

**DOI:** 10.1101/2024.10.10.617522

**Authors:** Yuchen Zhang, Wenkai Lai, Meiling Wang, Shirong Lai, Qing Liu, Qi Luo, Lijun Dou, Fenglong Yang

## Abstract

**Background:** Inflammatory bowel disease (IBD), encompassing ulcerative colitis (UC) and Crohn’s disease (CD), represents a chronic inflammatory condition with an incompletely understood etiology. Emerging evidence suggests that alterations in gut microbiota composition play a pivotal role in disease development. Here, we leveraged gut fecal metagenomic data (3044 samples: 2248 IBD and 796 healthy) from publicly available sources to explore microbiome biomarkers related to IBD to provide new ideas for clinical treatment.

**Results:** Our analyses revealed marked disparities in microbial species abundance and composition between IBD and healthy samples in both male and female subjects from the United States, whereas such distinctions were absent in subjects from Spain. Hierarchical and systematic investigations uncovered microbial phyla, such as Verrucomicrobia and Firmicutes, associated with IBD in US cohorts. Furthermore, we identified 127 highly correlated pathways with these differential microbes, covering functions such as peptidoglycan biosynthesis III, dTDP-L-rhamnose biosynthesis I, starch degradation, and glucose-1-phosphate degradation. Gut microbial metabolites were predicted based on metagenomic data using the MelonnPan workflow and 16 metabolites with significant differences were identified that collectively contribute to energy metabolism, digestion, skin integrity and general bodily function.

**Conclusions:** This study identified differences in microbial species and metabolic pathways related to Inflammatory Bowel Disease (IBD) through hierarchical and systematic analysis, which can aid in the clinical diagnosis of IBD. Pathways such as Peptidoglycan Biosynthesis III and dTDP-L-Rhamnose Biosynthesis I are associated with the generation of bacterial cell walls, and disruptions in these pathways affect bacterial activity, leading to an imbalance in the host’s gut microbiota. Based on the hypotheses regarding the pathogenesis of IBD derived from the above mining results, it provides a theoretical basis for selecting precise treatment options.

## Introduction

Inflammatory bowel disease (IBD) is a typical chronic inflammation that includes ulcerative colitis (UC) and Crohn’s disease (CD). The etiology of IBD remains unclear due to the complex interplay between genetic variability, host immune system, and environmental factors in this heterogeneous disease[1].Previous studies have consistently found a close association between dysbiosis of the gut microbiota and IBD. Research has shown that specific species within the host microbiota can regulate the host immune system, indicating the important role of specific bacteria at specific developmental stages in maintaining normal immune function[2], in a study comparing germ-free (GF) and specific pathogen-free (SPF) mice, found a significant induction of Treg cells in the intestines of SPF mice compared to GF mice. This induction was associated with the presence of *Clostridium* species in the mouse gut[3].

Changes in the composition of the gut microbiota can lead to alterations in metabolites, which may play a role in the pathogenesis of IBD. Through multi-omics correlation analysis (metagenomics and metabolomics), it is possible to establish correlations between the microbial composition and specific bacterial metabolic pathways and assess the impact of small-molecule metabolites on the mechanisms underlying IBD[4, 5]. When comparing the gut microbiota of healthy individuals and IBD patients, it was found that 12% of metabolic pathways showed significant differences, while only 2% of taxonomic species abundance at the genus level exhibited significant differences. Specifically, in the IBD microbiota, there was a decrease in amino acid biosynthesis and carbohydrate metabolism pathways, which may favor nutrient absorption and secretion processes. Additionally, the expression of genes associated with oxidative stress, such as glutathione and sulfate transporters, was also increased[5, 6].These findings may reflect the microbial response to the intestinal environment in inflammatory bowel disease (IBD) and suggest that, when studying microbial dysbiosis, functional differences at the functional level may be more compelling than species composition differences.

The gut microbiota is a critical regulator of gastrointestinal digestion, playing an important role in the extraction, synthesis, and absorption of various nutrients and metabolites[7]. Additionally, the gut microbiota has immunological functions by inhibiting the growth of pathogens[8], consuming nutrients, and producing bacteriocins to counteract pathogenic colonization in the gastrointestinal tract. Therefore, in this experiment, publicly available macrogenomic data, including functional expression profiles and relative abundance data of species, were used. Through stratified analysis based on sample metadata and systematic analysis integrating predicted metabolomic results, the experiment aimed to deeply explore multidimensional microbial biomarkers of IBD. Firstly, feature species were selected based on their occurrence frequencies in IBD and healthy samples, followed by hierarchical clustering analysis and stratification of samples based on the results and sample metadata. Subsequently, gut microbiota diversity analysis, species composition analysis, and differential analysis were performed to identify significantly different species[9, 10, 11]. Then, through association analysis between the microbiome and host phenotypes, pathways highly correlated with differentially abundant species were identified.

## Data

### Data Sources

The data used in this experiment were obtained from the curatedMetagenomicData package[12]. Figure 1 provides specific details about the data.

**Fig. 1.**
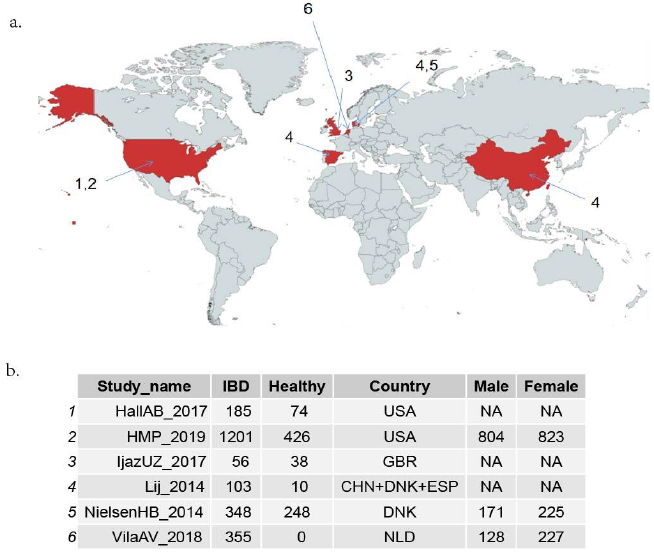
a.the distribution of samples; b. The details of data. All the raw metagenomic sequencing data underwent standardized upstream analysis using a unified pipeline. Bacterial abundance annotation was performed using the MetaPhlAn2 pipeline, while functional analysis was conducted using the HUMAnN2 pipeline. The phenotypic information of all datasets was reannotated and redefined using a standardized set of criteria. Finally, each dataset was packaged as an ExpressionSet object.

Due to uneven distribution of clinical information in the samples used in this experiment, samples were filtered to include only those that simultaneously had sample IDs, gender, age, nationality, and BMI index information. This resulted in 625 samples, including 352 IBD samples and 273 healthy samples. However, among these 625 samples, all 121 samples from Denmark were healthy samples, and there were no IBD samples from the same nationality available for comparison. Therefore, these samples were excluded from subsequent differential analysis. Consequently, the final dataset comprised 497 samples, including 347 IBD samples and 150 healthy samples, which were used for further differential analysis (Figure 2).

**Fig. 2.**
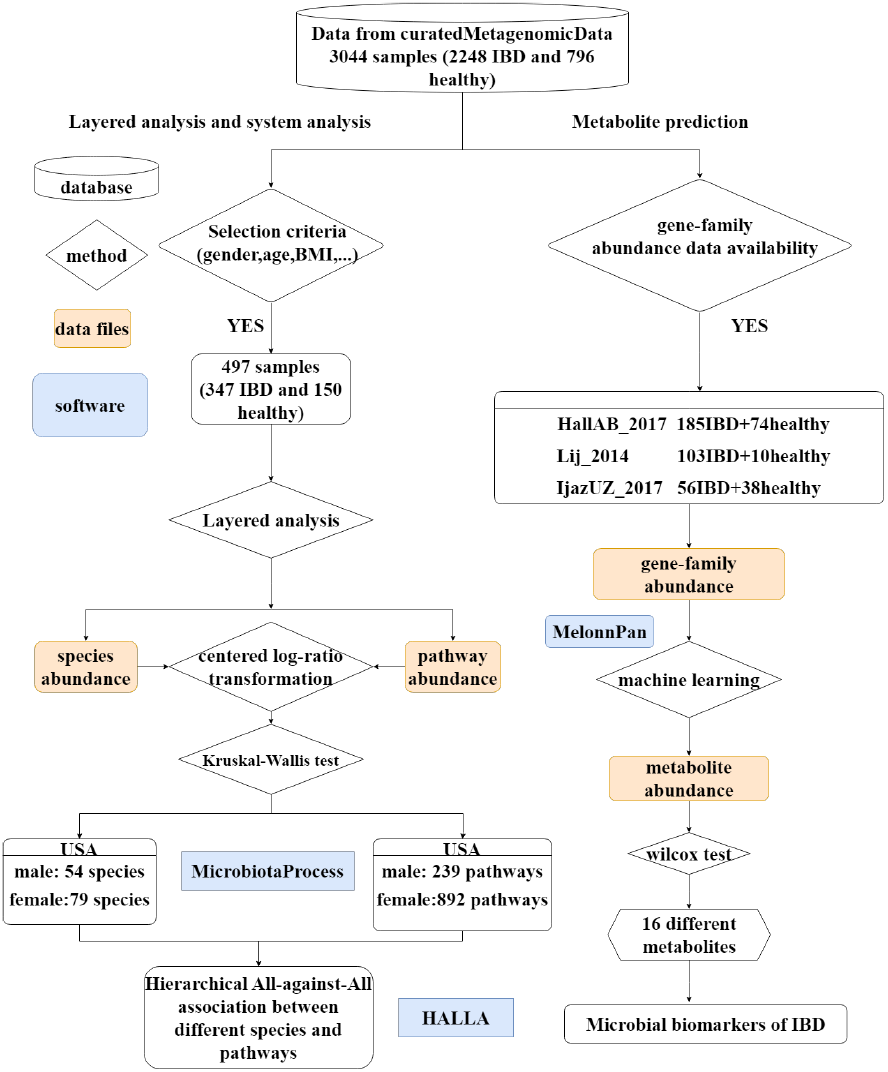
The flow chart of this project

#### Data processing of species expression profiles

The relative abundance data of the microbiota in the samples were obtained from a public database. The relative abundance data of the microbiota are compositional data, reflecting the proportions of each bacterial species within the total bacterial community in each sample. The characteristic feature of compositional data is that the sum of the relative abundances of all bacterial species in each sample is equal to 1. Therefore, preprocessing of the relative abundance data was performed by excluding bacterial species with relative abundances of 0 in more than 90% of the samples, resulting in 171 bacterial species. Subsequently, the frequency of occurrence of these species in IBD samples (a) and healthy samples (b) was calculated, and a species filtering index was constructed as index = a/b. For each sample, the index was calculated.

Following that, a centered log-ratio transformation was applied to the remaining relative abundance data of the microbiota.

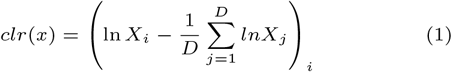

#### Data processing of pathways expression profiles

The functional expression profiles used in this experiment were obtained by establishing a linear regression model between the metabolite annotation results and the measured concentrations of metabolites in the reference database. This allowed for quantitative analysis of the metabolites in the samples. Subsequently, the metabolites were mapped to metabolic pathways in the reference database to determine their functional roles and involvement in biological processes, resulting in the generation of functional expression profiles.The abundance of gene families was calculated as the weighted sum of alignments obtained from each read, and normalization was performed based on gene length and alignment quality.

## Methods

### Mining scheme for gut microbiota biomarkers

Firstly, download the relative abundance, metabolic pathway expression profile, genome and other data of the samples from the database, screen the samples, calculate the alpha and beta diversity of the samples, and use genome data for metabolite prediction. Then, perform differential analysis on the relative abundance, metabolic pathway, and metabolite abundance of the samples, and perform correlation analysis on the differential microbiota and differential pathway to obtain biomarkers related to IBD(Fig.2).

### Calumniate the Gut Microbiome Health Index(GMHI)

There were significant differences in age and BMI index between the IBD samples and healthy samples, therefore, we used the Gut Microbiome Health Index (GMHI) to reflect the human gut health status. The GMHI is a measure proposed in a study published in Nat Commun in 2020, which evaluates health status based on species-level gut microbiota composition[13]. It is calculated using 50 specific microbial species. The calculation formula for GMHI is as follows:

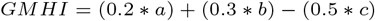

Whereas, a refers to the relative abundance of beneficial microbiota in the human body; b refers to the relative abundance of commensal microbiota coexisting with the human body; c refers to the relative abundance of detrimental microbiota in the human body. According to the literature, samples with GMHI more than 0 are considered healthy samples, while samples with GMHI less than 0 are considered disease samples. The index achieved an accuracy of 73.7%. Therefore, for the 497 samples included in the experiment, the GMHI was calculated for each sample, and the classification performance was evaluated.

### Microbiota alpha Diversity analysis

Alpha diversity refers to the biodiversity within a single sample and is often used to describe the richness and evenness of species in an ecosystem. In the context of microbial communities, alpha diversity is commonly used to assess the number of microbial species and the evenness of their relative abundance distribution within a sample. Alpha diversity is calculated using indices based on OTUs (Operational Taxonomic Units) or ASVs (Amplicon Sequence Variants), such as Shannon index, Simpson index, Chao1 index, etc. The level of alpha diversity reflects the complexity and stability of the microbial community. A high alpha diversity indicates high species richness, even relative abundance distribution, and better stability of the microbial community within a sample. Conversely, a low alpha diversity suggests low species richness, uneven relative abundance distribution, and poorer stability of the microbial community.

In this experiment, alpha diversity, including indices such as Chao1, Shannon, and Simpson, was calculated. The non-parametric Kruskal-Wallis rank-sum test was used to assess the differences in alpha diversity among different conditions or states.

### Beta Diversity analysis and sample clustering analysis

Beta diversity refers to the dissimilarity in species composition between different ecosystems or groups. It captures the differences in species composition among different groups and reflects the differentiation between biological communities. Beta diversity primarily utilizes the species composition or evolutionary relationships and abundance information among samples to calculate distances between samples, aiming to identify significant differences in microbial communities among samples. Commonly used methods for beta diversity analysis include Principal Component Analysis (PCA) and Principal Coordinates Analysis (PCoA). PCA is a widely used method for data dimensionality reduction and visualization. It transforms high-dimensional data (with multiple feature variables) into low-dimensional data (with fewer feature variables) while retaining most of the original data’s information and variance. PCA involves standardizing each feature variable to have a mean of 0 and a variance of 1, calculating the covariance matrix to determine the correlation between feature variables, computing the eigenvalues and eigenvectors of the covariance matrix, and selecting the top k eigenvectors based on the eigenvalues’ magnitude as principal components. PCoA has a similar principle to PCA but uses a distance matrix between samples instead of a correlation matrix.

Hellinger transformation involves taking the square root of each element in the relative abundance matrix and normalizing all elements within each sample to sum up to 1. This transformation reduces sparsity and skewness and provides a better reflection of microbial community diversity. In this study, the relative abundance data was Hellinger transformed, and PCoA based on the Bray-Curtis distance was used for beta diversity analysis.

Hierarchical clustering is a clustering method based on distance measurement. It iteratively merges the most similar samples or subgroups to form a hierarchical tree-like structure. In the clustering tree, the original data points of different categories are at the lowest level, and the top level represents the root node of a cluster. Different distance measurement methods can be used to calculate the distances between samples, such as Euclidean distance, Manhattan distance, and Chebyshev distance. In this experiment, the Euclidean distance was used as the distance measurement method for hierarchical clustering.

### Analysis of Differences in Microbial Composition and Function

All species composition analysis and diversity analysis were performed in R software using the MicrobiotaProcess and Maaslin analysis pipelines. MicrobiotaProcess defines the MPSE (MicrobiotaProcess Sequence Experiment) data structure, which integrates various formats of upstream microbiome data and downstream analysis outputs. It utilizes the tidy framework to develop analysis functions such as data filtering and normalization, phylogenetic transformation, identification and visualization of differential species, and provides a unified and concise analysis syntax for downstream analysis of microbiome data. This framework facilitates the management and reproducible analysis of microbiome data[9].

### Correlation analysis between gut microbiota and host phenotype

This study employed the MaAsLin2 (Microbiome Multivariable Association with Linear Models) method to investigate the relationship between gut microbiota and host phenotypes. MaAsLin2 is a multivariate linear regression model specifically designed for microbiome data, published in PLoS Comput Biol in 2021[14]. Microbiome data is characterized by high dimensionality and sparsity, with many microbial taxa only present in a subset of samples and a large number of zeros. In extreme cases, the abundance of individual taxa in the control group can be zero. Therefore, MaAsLin2 preprocesses the microbiome data, including data normalization, to address these challenges. Normalization aims to compare the microbial features among different samples, irrespective of their absolute values. Common normalization methods include log2 transformation and total sum normalization. After preprocessing the data, MaAsLin2 employs a multivariate linear regression model to model the relationship between microbial features and environmental/clinical features. In MaAsLin2, the model can be formulated as follows:

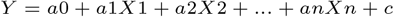

Where Y represents the environmental or clinical feature (dependent variable), X1, X2, …, Xn represent the microbial features (independent variables), X0, X1, X2, …, Xn represent the coefficients of the model, and c represents the error term. The coefficients X in the model can explain the relationship between microbial features and environmental/clinical features. For example, the coefficient X1 indicates that, holding other factors constant, a one-unit change in X1 will result in a X1 unit change in Y.

### Correlation analysis of microbial community structure and function

This study utilized the HAllA (Hierarchical All-against-All association) method to perform association analysis between differential species and sample functional profiles. HAllA is a method published in BioData Min in 2017 that aims to identify correlated feature blocks within high-dimensional datasets[15].The specific steps are as follows: Firstly, the microbiome data is normalized. Then, the HAllA method is used to explore the associations between microbiome features.

In detail, HAllA divides the microbiome features into multiple groups, each containing similar microbiome features. For each group, HAllA calculates the associations between the features and selects the most significant association as the representative association for the group. This approach reduces the dimensionality of the microbiome features from a high-dimensional space to a low-dimensional space, thus reducing computational complexity. Next, a joint test statistic is used to determine which microbiome features are associated with the environmental or clinical features. Specifically, for each group, HAllA calculates the associations between all microbiome features within the group and the environmental or clinical features, and selects the most significant association as the representative association for the group. The similarity between two sets of data is measured using the Spearman rank correlation, and the resulting p-values are corrected using the Benjamini-Hochberg false discovery rate (FDR) correction. If the association of a microbiome feature exceeds a threshold, it is considered to be associated with the environmental or clinical feature.

### Prediction of gut microbiota metabolites

Metabolites are the end products of microbial metagenomics and play a crucial role in disease progression. Therefore, in this experiment, the MelonnPan workflow is used to predict metabolites associated with microbial communities using the metagenomic data of the samples. MelonnPan is a predictive model that infers microbial community metabolite features from amplicon or metagenomic data. The MelonnPan model can be trained to predict metabolite profiles for specific microbial community types. The training dataset consists of paired metagenomic and metabolomic data from the target environment. It utilizes elastic net regularization regression for each metabolite to identify the minimal microbial feature set that predicts its abundance. Cross-validation is used to assess these models and identify metabolites that are not well-suited for prediction. The result is a coefficient matrix for the sequence of metabolite features. This matrix can be applied to new metagenomic data to predict the corresponding metabolite features.

## Result

### Stratified Analysis

#### Based on the abundance-based sample clustering, a hierarchical pattern was observed in relation to nationality and gender

For the 497 samples, the occurrence frequencies (a) of all species in the IBD samples and the occurrence frequencies (b) in the healthy samples were calculated. An index was constructed as a/b. Feature species were selected in each group based on the index and subjected to hierarchical clustering, as shown in the figure. The heatmap indicates a potential association between the presence of IBD and nationality and gender. To further investigate, all samples were grouped based on the clustering analysis results and sample metadata, specifically nationality and gender, for subsequent analysis.

The significance of the ANOSIM test, represented by the R value, indicates the degree of dissimilarity between groups. When R tends to 1, it indicates that the differences between groups are greater than the differences within groups. When R = 0, it indicates that there are no differences between groups. When R tends to -1, it indicates that the differences between groups are smaller than the differences within groups. After conducting ANOSIM tests on the stratified samples, except for Spanish female samples, the R values were greater than those of the combined samples before stratification. Therefore, subsequent differential analyses were based on this result.(Fig.3)

**Fig. 3.**
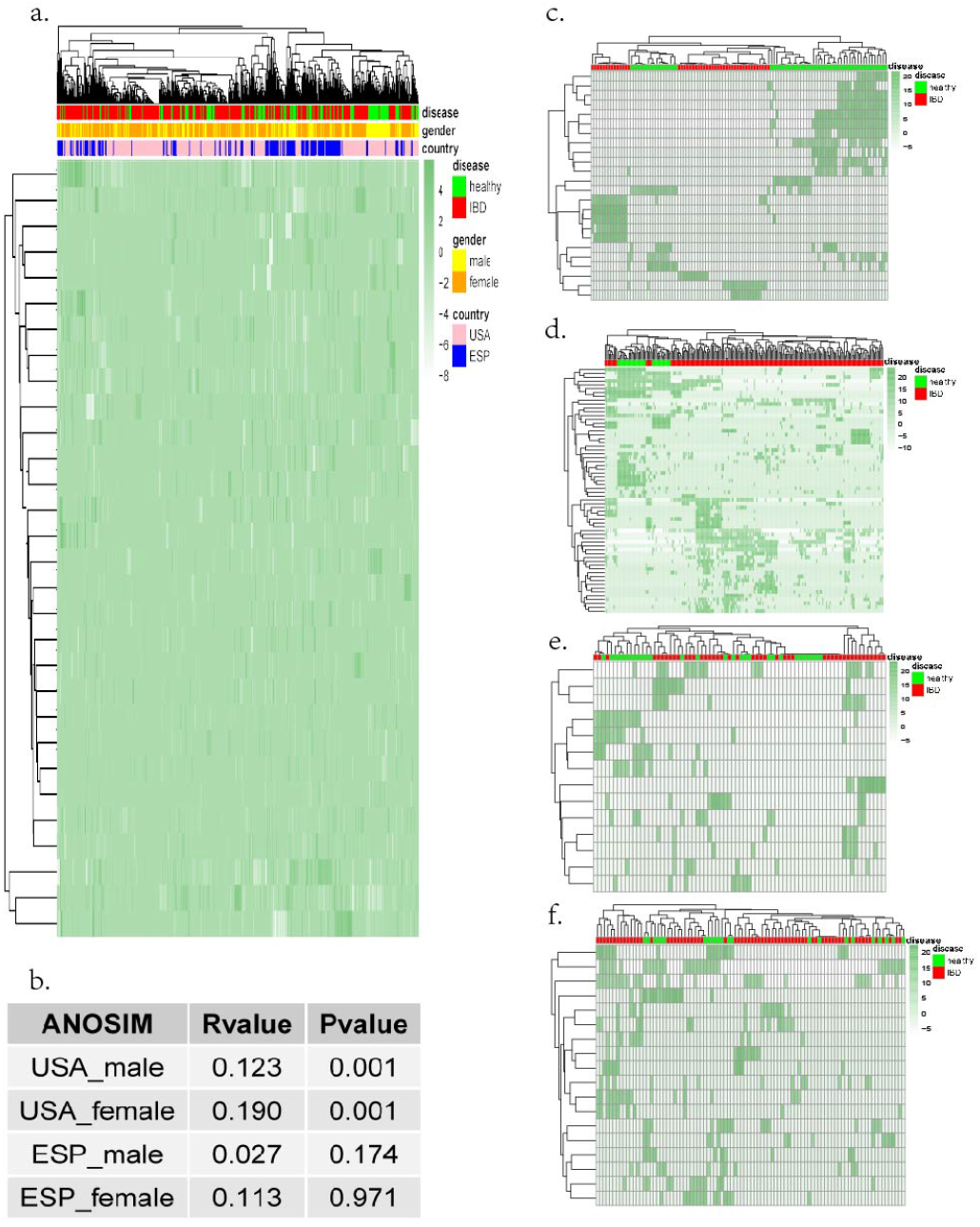
a.Hierarchical clustering of sample similarity based on relative abundance of microbial communities of all samples,ANOSIM R value:-0.04682,Significance: 0.99 b.the table of ANOSIM’s statistics c.Hierarchical clustering of sample similarity based on relative abundance of microbial communities of USA male samples d.Hierarchical clustering of sample similarity based on relative abundance of microbial communities of USA female samples e.Hierarchical clustering of sample similarity based on relative abundance of microbial communities of ESP male samples f.Hierarchical clustering of sample similarity based on relative abundance of microbial communities of ESP female samples

#### Species of gut microbiota alpha Diversity stratification analysis

Alpha diversity index reflects the number of species and their relative abundances within a community, which is the result of competition or symbiosis among species utilizing the same habitat. Comparing the alpha diversity indices of different samples can reveal differences in diversity. In this experiment, the Chao index, Shannon index, Simpson index, and Pielou index were calculated for disease samples and normal samples in the four groups of data to reflect the differences between disease and normal samples. The results of the alpha diversity analysis showed significant differences in species abundance and species composition between disease and normal samples, except for the Pielou index in the American male samples.

The relative abundances of species in the four previously grouped datasets were standardized, followed by PCoA analysis. PCoA analysis showed good classification performance in American male and female samples. Based on the results of alpha diversity analysis and PCoA analysis, there were no significant differences in the gut microbiota between IBD and normal populations in the two Spanish sample groups. Therefore, subsequent differential analysis mainly focused on the American male and female samples(Fig.S1-Fig.S4).

#### Species composition and differential stratification analysis of gut microbiota

It is well known that the human gut microbiota is mainly composed of five major phyla: Firmicutes, Bacteroidetes, Proteobacteria, Actinobacteria, and Fusobacteria. The Fig.S5 shows the top 5 most abundant species in USA male samples(left) and USA female samples(right) and performs hierarchical clustering of the samples.

Through species composition analysis and differential analysis, this study found that in the US male samples, Verrucomicrobia was only found in healthy samples. In addition, the abundance of Firmicutes was higher in most healthy samples compared to IBD samples. In the US female samples, a similar distribution pattern of Firmicutes abundance was observed. However, in the Spanish male and Spanish female samples, there were no significant differences in the composition of gut microbiota between IBD and healthy samples. It is speculated that in these two datasets, IBD occurrence may not be caused by gut microbiota. The Kruskal-Wallis test and Wilcoxon rank-sum test were subsequently used to perform differential analysis on the four datasets. Box plots were used to display the relative abundance of differentially abundant species after sorting. The Kruskal-Wallis test is a non-parametric method used to determine if two or more samples come from the same probability distribution. Unlike one-way ANOVA, the K-W test does not assume that the samples come from a normal distribution. The following figure shows the distribution of differentially abundant species in the two datasets. The left side of the figure displays box plots of the relative abundance of differentially abundant species in the disease and healthy samples, while the right side displays the sorted lg(LDA) values of the differentially abundant species. The box plots reveal that there is a significant difference in the types of differentially abundant species between the US male and US female samples. However, some species, such as Agathobaculum butyriciproducens, exhibit significant differences in both datasets(Fig.4).

**Fig. 4.**
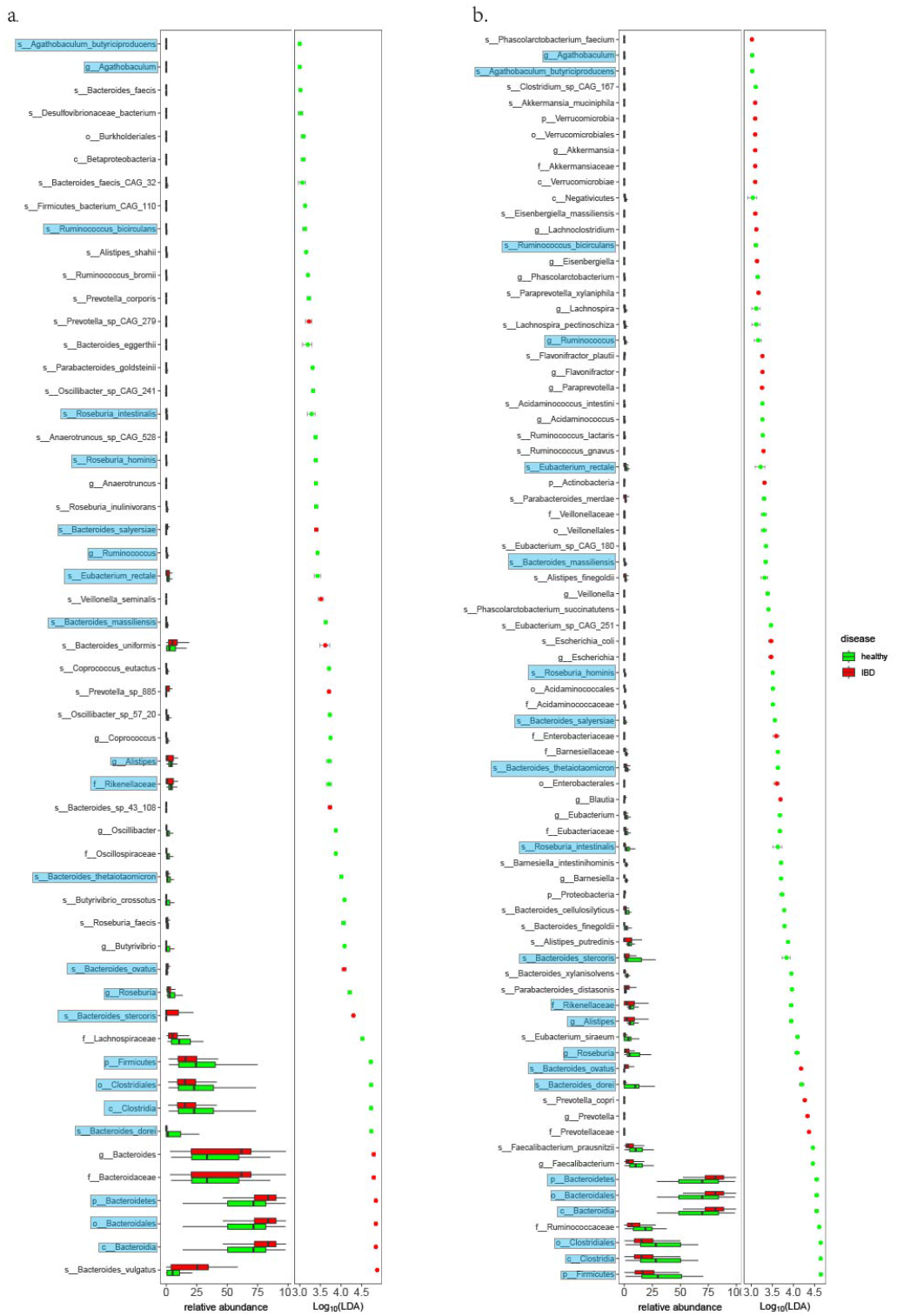
a.The results of differential analysis of male samples in the United States and the relative abundance of various bacterial species are sorted according to the lg (LDA) value b.The results of differential analysis of female samples in the United States and the relative abundance of various bacterial species are sorted according to the lg (LDA) value

Agathobaculum butyriciproducens is a Gram-positive, obligate anaerobic bacterium. Although it is present at low levels in healthy individuals, it is significantly reduced in patients with inflammatory bowel disease and other intestinal disorders. This suggests that Agathobaculum butyriciproducens may play a beneficial role in maintaining gut health.

#### Performing species differential analysis using a multivariable linear regression model with GMHI as a covariate

Preliminary exploration of the data revealed significant differences in age and BMI index between IBD patients and healthy individuals. Additionally, previous studies have shown that GMHI can effectively assess human gut health. Therefore, for the 497 samples included in the study, the GMHI values were calculated for each group, and scatter plots were generated. Samples with GMHI values greater than 0 were considered healthy, while samples with GMHI values less than 0 were considered diseased. Attempts were made to match the samples based on age, BMI index, and GMHI using propensity score matching. However, the matched sample size was too small to meet the requirements for subsequent analyses.

Due to the significant heterogeneity in gut microbiota composition among individuals, the GMHI calculated based on specific species was unable to distinguish between healthy and IBD samples. In order to further explore gut microbiota markers for IBD, the GMHI values were modified as follows: if the GMHI value of a sample was less than 0, it was replaced with 0; otherwise, it was replaced with 1. These modified GMHI values were then used as covariates in the analysis using MaAsLin2. In the case of the American male samples, the differential species Bacteroides vulgatus and Bacteroides dorei identified by MaAsLin2 analysis were also found in the previous results.

Bacteroides vulgatus exhibits significantly higher abundance in IBD samples compared to normal samples. Bacteroides vulgatus is a common Gram-negative anaerobic bacterium found in the human gut. It belongs to the Bacteroides genus and is one of the most abundant and diverse bacterial populations in the human gut microbiota. Bacteroides vulgatus is capable of breaking down complex carbohydrates, such as dietary fibers, and fermenting them into short-chain fatty acids (SCFAs). The SCFAs produced by Bacteroides vulgatus and other gut bacteria play important roles in maintaining a healthy gut microbiota and overall health. SCFAs provide energy to the cells lining the colon, help maintain intestinal barrier integrity, and prevent inflammation. They also act as signaling molecules, regulating systemic immune function and metabolism. Bacteroides vulgatus has been associated with various human diseases, including inflammatory bowel disease, colon cancer, and obesity. For instance, when it overgrows in the gut, it can lead to inflammation and other health issues. Studies, such as the one conducted by Li et al., have shown that Bacteroides vulgatus overgrowth in the gut of individuals with coronary artery disease is closely associated with inflammatory and metabolic dysregulation processes[11].

Bacteroides dorei exhibits significantly lower abundance in IBD samples compared to normal samples. Bacteroides dorei is a species of bacteria belonging to the Bacteroides genus, a common group of gut bacteria. It is specifically named Bacteroides dorei. Bacteroides dorei plays a role in the breakdown and absorption of complex carbohydrates, proteins, and fats from food, thereby contributing to energy supply in the human body. Studies have suggested that a deficiency of Bacteroides dorei is associated with the occurrence and progression of certain diseases, such as chronic intestinal inflammation and colorectal tumors[16]. Through MaAslin2 analysis, only five significantly different bacterial species were found in the US female samples, and these five different species did not overlap with the previous experimental results.

#### Stratified analysis of differences in gut microbiota metabolic pathways

First, hierarchical clustering was performed on the relative abundance of metabolic pathways in all samples, followed by ANOSIM testing.

Hierarchical clustering heatmaps and ANOSIM test results showed that it was not possible to distinguish between IBD samples and normal samples. In order to further investigate the differences in metabolic pathways between IBD and normal samples, the samples were stratified based on nationality and gender, and hierarchical clustering and ANOSIM tests were performed on the stratified groups.

The ANOSIM test conducted on the stratified samples showed that the R values for both groups were significantly higher than those for the combined samples before stratification. Therefore, subsequent analyses were based on this stratified grouping(Fig.5).

**Fig. 5.**
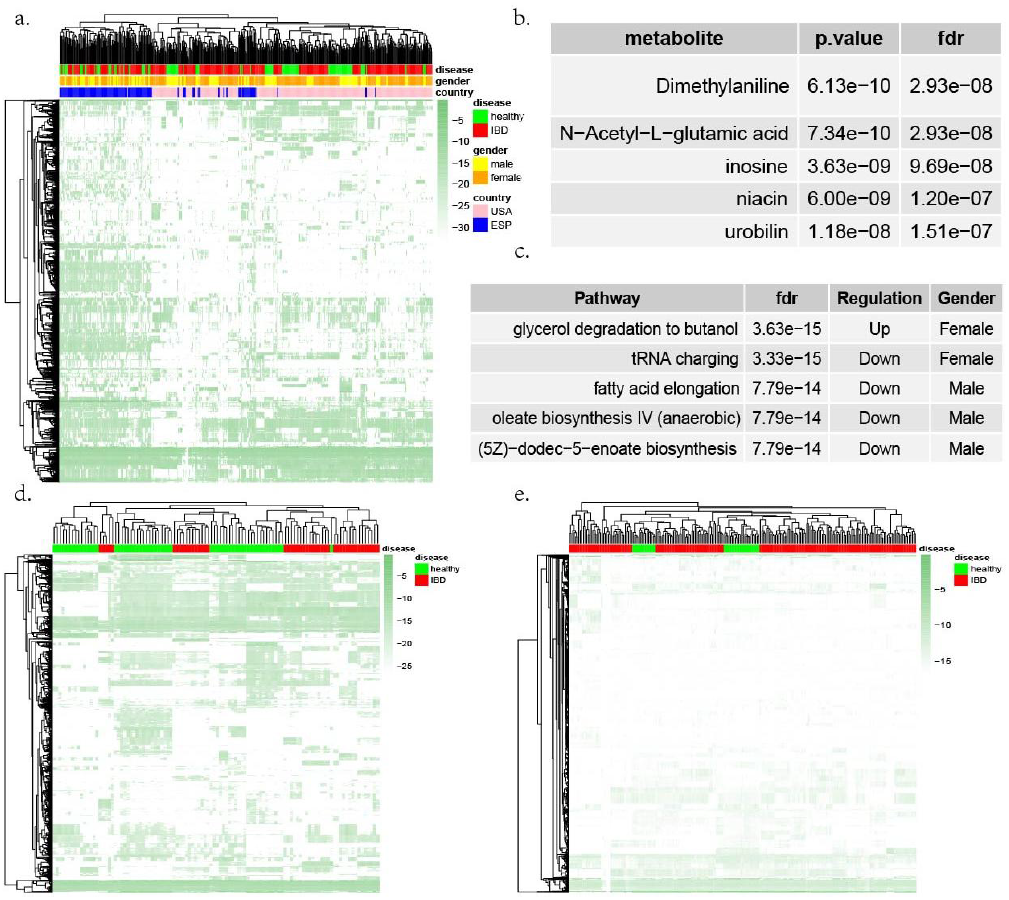
a.Hierarchical clustering of sample similarity based on pathway abundance of all samples b.The table of different metabolites. c. The table of different pathways d.Hierarchical clustering of sample similarity based on pathway abundance of USA male samples e.Hierarchical clustering of sample similarity based on pathway abundance of USA female samples

First, the Shapiro-Wilks test was used to assess the normality of the data, and it was found that the data did not follow a normal distribution. Hence, the Wilcoxon rank-sum test was employed to assess the differences between IBD samples and normal samples, and the p-values were adjusted using the Benjamini-Hochberg method. A significance threshold of FDR ¡ 0.00005 was applied, resulting in the identification of 599 pathways with significant differences. These pathways included TRNA-CHARGING-PWY: tRNA charging, FASYN-ELONG-PWY: fatty acid elongation – saturated, and others.

The differential analysis results indicate that the dysregulated pathways in the gut microbiota of IBD patients are mainly associated with important biological processes such as fatty acid synthesis and protein synthesis. Disruptions in these pathways can impact the metabolic products of the gut microbiota, disrupt the host’s gut microbiota balance, and contribute to the development of IBD.

### Meta-analysis

#### Prediction of gut microbiota metabolites

The gut microbiota ultimately affects the host through its own metabolites. Therefore, in this experiment, metagenomic data was used with machine learning methods to predict metabolites. Each sample obtained data on 80 metabolites. The Wilcoxon rank-sum test was used to perform differential analysis on the 80 metabolites, and the p-values were corrected using the Benjamini-Hochberg method. After screening for FDR (false discovery rate) less than 1e-7, 30 metabolites with significant differences were identified. The top 5 metabolites are shown in Fig5e(Fig.5).

The 16 different metabolites have diverse roles in the human body. They include fatty acids like adrenic acid, bile acids such as chenodeoxycholate and deoxycholic acid aiding fat digestion, urobilin involved in urine color, and compounds like ceramides for skin health. N-Acetylglutamate and nicotinic acid are essential for metabolic processes. Amino acid derivatives like dimethyllysine and ADMA impact protein function and blood vessels. Inosine is a nucleoside with various roles, and bilirubin reflects liver and gallbladder health. These metabolites collectively contribute to energy metabolism, digestion, skin integrity, and overall bodily function.

#### Correlation analysis between species and functional pathways

We use Spearman’s rank correlation as a measure of similarity between the relative abundance of gut microbiota and the significant pathways. The resulting p-values were corrected using the Benjamini-Hochberg false discovery rate (FDR) correction. This analysis aimed to identify highly correlated pathways with the relative abundance of differential microbial species.In this section, we will conduct correlation analysis on the differential species and differential functional pathways between IBD samples and normal samples in two sets of data(Fig.S5-Fig.S8).

For example, in the male samples from the United States, we identified a positive correlation (0.447) between Alistipes indistinctus and PWY-7219: adenosine ribonucleotides de novo biosynthesis(Bacteroides vulgatus CAG 6) in healthy samples, while the correlation in IBD samples was nearly zero (−0.082). Similarly, in female samples from the United States,we found a negative correlation (−0.7018) between Bacteroides thetaiotaomicronand PWY-6124: inosine-5’-phosphate biosynthesis II in healthy samples, whereas it was positive correlated (0.1238) in IBD samples(Fig.S9).As is well known, there is a close relationship between gut microbiota and gut health. Gut microbiota participate in food digestion and breakdown, particularly in the degradation of complex carbohydrates and cellulose. They produce beneficial metabolites such as short-chain fatty acids, which contribute to maintaining gut mucosal health and nutrient absorption. Additionally, gut microbiota are intricately linked to the immune system. They help regulate immune responses, maintain immune balance, prevent excessive immune activation and inflammatory responses, thereby reducing the risks of autoimmune diseases and allergies.Based on these results, we speculate that:(1)In the process of IBD development, the strength of interconnections among various species within the gut microbiota decreases,contributing to the progression of the disease. (2)During the occurrence of IBD, certain species’ metabolic pathways are disrupted, and the metabolites generated from these abnormalities affect the host gut environment, accelerating the disease process.

## Discussion

The human microbiota is a diverse microbial ecosystem that is associated with numerous beneficial physiological functions as well as the pathogenesis of many diseases. The microbiota is primarily composed of bacteria but also includes fungi, viruses, archaea, and protists, forming a symbiotic community. Trillions of microorganisms colonize the human body, forming the microbiota, which is collectively referred to as the human microbiome. The human microbiome spans different domains, including bacteria, eukaryotes, archaea, and viruses, and plays crucial roles in energy acquisition, immune development, and providing essential protection, structural support, and metabolic functions that are vital to human physiology and health maintenance[17].The advancements in next-generation sequencing and computational technologies have provided opportunities for studying the structure and function of microbial communities associated with various body sites.

The gut microbiome is an ecosystem composed of various microorganisms including bacteria, fungi, archaea, and viruses. Under normal conditions, the gut microbiome maintains intestinal balance in conjunction with the host and participates in numerous physiological processes such as energy metabolism, nutrient absorption, and immune regulation. However, in patients with inflammatory bowel disease (IBD), the composition and function of the gut microbiome undergo alterations that may lead to impaired intestinal barrier, dysregulation of the immune system, and exacerbated inflammation. Several studies have demonstrated significant differences in the gut microbiome between IBD patients and healthy individuals. For instance, there are commonly observed increases or decreases in certain genera within the gut microbiome of IBD patients, such as Clostridium, Bacteroides, and Bacillus. Moreover, the diversity and stability of the gut microbiome are often reduced in IBD patients compared to healthy individuals.

The relationship between the gut microbiome and IBD is bidirectional. On one hand, dysbiosis of the gut microbiome may contribute to the development and progression of IBD. On the other hand, the inflammatory response and medications used to treat IBD can also influence the composition and function of the gut microbiome. Therefore, modulation of the gut microbiome has emerged as a potential strategy for the prevention and treatment of IBD.

Current evidence suggests that interventions targeting the gut microbiome, such as the use of probiotics, prebiotics, and fecal microbiota transplantation, can improve clinical symptoms and reduce intestinal inflammation in IBD patients. However, the mechanisms underlying the role of the gut microbiome in IBD are still not fully understood and require further investigation.

At the metabolomic level, IBD patients may exhibit alterations in the abundance and proportions of common metabolites such as amino acids, lipids, and carbohydrates. Additionally, IBD can affect metabolic pathways and networks, including glycolysis, fatty acid metabolism, and amino acid metabolism. Studies have identified characteristic changes in the metabolomic profiles of IBD patients, such as accumulation of glycolytic products and reduction of fatty acid metabolites, which may be associated with inflammation and immune dysregulation in IBD.

The gastrointestinal tract is the most densely populated microbial habitat in the human body. Despite the recognized beneficial functions of the gut microbiota, it is associated with various pathological conditions, including inflammatory bowel disease (IBD) and its two main entities: Crohn’s disease (CD) and ulcerative colitis (UC). Numerous studies have demonstrated that the chronic inflammation of the intestinal mucosa in IBD is associated with alterations in the structure and function of the microbial community, known as “dysbiosis.” Dysbiotic bacterial microbiota has been extensively characterized in IBD[18, 19],However, the roles of other microbial components and their interactions within the gut microbiota are still not fully understood.

In this experiment, publicly available microbiome data were collected, and the samples were subjected to stratified analysis and systematic analysis. Analysis pipelines such as MicrobiotaProcess were employed to analyze the data. Several bacterial species, including Verrucomicrobia and Firmicutes, were found to be associated with IBD. Correlation analysis revealed a strong association between differential bacterial species and pathways such as PWY-6168: flavin biosynthesis III, PWY-7371: 1, PWY-621: sucrose degradation III (sucrose invertase), and PWY-7242: D-fructuronate degradation.

Verrucomicrobia is a bacterial phylum widely present in natural environments, and in the human gut, it enhances the integrity and stability of the intestinal mucosa by producing specific polysaccharides. This protection helps prevent harmful microorganisms and toxins from damaging the intestine. Additionally, Verrucomicrobia can maintain intestinal homeostasis by modulating immune responses, thereby preventing inflammation and autoimmune diseases.

Certain bacteria within Firmicutes have the ability to break down polysaccharides and fatty acids, producing organic acids and gases as metabolic byproducts. These byproducts serve as nutrients for other gut bacteria and also influence physiological processes such as intestinal pH, oxygen levels, and intestinal motility. Firmicutes are also involved in polysaccharide and fatty acid metabolism, obesity and metabolic-related diseases, immune regulation, and maintenance of the intestinal mucosal barrier.

Overall, these findings highlight the potential involvement of Verrucomicrobia and Firmicutes, along with their associated pathways, in the pathogenesis and development of IBD. Further research is needed to elucidate the precise mechanisms and interactions within the gut microbiota that contribute to IBD and to explore their therapeutic implications.

However, this experiment has several limitations. Firstly, the samples only included populations from the United States and Spain, which may introduce geographic biases in the analysis results. It is necessary to collect more diverse samples for future studies. Secondly, the 127 pathways highly correlated with differential bacterial species obtained from the correlation analysis lack further exploration and validation. More in-depth investigations are required to understand the functional implications of these pathways in IBD.

## Conclusion

Based on the stratified analysis and integrated analysis, this experiment identified Verrucomicrobia, Firmicutes, and other taxa that may be associated with IBD. Additionally, we found high correlations between the differential bacterial species and pathways such as PWY-6168: flavin biosynthesis III, PWY-7371: 1, PWY-621: sucrose degradation III (sucrose invertase), and PWY-7242: D-fructuronate degradation. We also found significant differences in the correlation between certain pathways and species between IBD samples and normal samples, which may be related to the process of IBD.

## Key Points

Sample clustering based on species abundance reveals the stratification phenomenon of nationality and gender.

The difference analysis results indicate that there are 22 significantly different species in both the male and female samples of the United States

The results of correlation analysis indicate that there are significant differences in the correlation coefficients between some species and functions between disease samples and healthy samples, such as Alistipes indistinctus and PWY-7219: adenosine ribonucleates de novo biosynthesis (Bacteroides vulgatus CAG 6); Eubacterium sp CAG 274 and PWY-5097: L-lysine biosynthesis VI (s Roseburia integralis)

## Supporting information

https://github.com/zyc134/IBD/archive/refs/heads/master.zip

## Supplementary Material

## Data Availability Statement

## Authors

