## Supplementary figures and images for "Systematic Mining of gut microbiota biomarkers for IBD"

### FigureS5.png

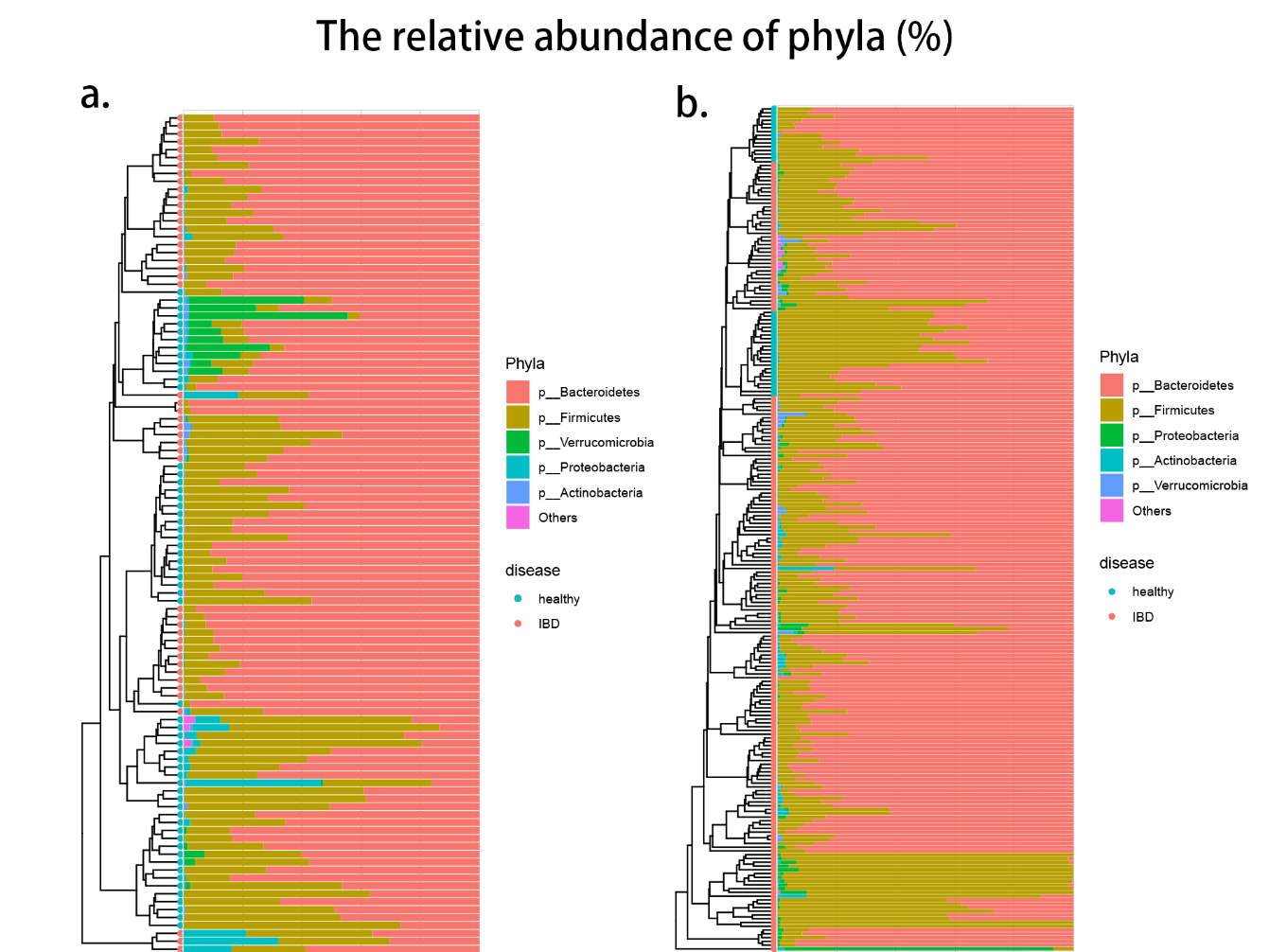
